# Cryo-EM structures of human arachidonate 12S-Lipoxygenase (12-LOX) bound to endogenous and exogenous inhibitors

**DOI:** 10.1101/2023.03.10.532002

**Authors:** Jesse I. Mobbs, Katrina A. Black, Michelle Tran, Hariprasad Venugopal, Theodore R. Holman, Michael Holinstat, David M. Thal, Alisa Glukhova

**Author notes:** These authors made equal contributions (Jesse I. Mobbs, Katrina A. Black).

## Abstract

Human 12-lipoxygenase (12-LOX) is an enzyme involved in platelet activation and is a promising target for antiplatelet therapies. Despite the clinical importance of 12-LOX, the exact mechanisms of how it affects platelet activation are unclear, and the lack of structural information has limited drug discovery efforts. In this study, we used single-particle cryoelectron microscopy to determine the high-resolution structures (1.7 Å - 2.8 Å) of human 12-LOX for the first time. Our results showed that 12-LOX can exist in multiple oligomeric states, from monomer to hexamer, which may impact its catalytic activity and membrane association. We also identified different conformations within a 12-LOX dimer, likely representing different time points in its catalytic cycle. Furthermore, we were able to identify small molecules bound to the 12-LOX structures. The active site of the 12-LOX tetramer is occupied by an endogenous 12-LOX inhibitor, a long-chain acyl-Coenzyme A. Additionally, we found that the 12-LOX hexamer can simultaneously bind to arachidonic acid and ML355, a selective 12-LOX inhibitor that has passed a phase I clinical trial for treating heparin-induced thrombocytopenia and has received fast-track designation by the FDA. Overall, our findings provide novel insights into the assembly of 12-LOX oligomers, its catalytic mechanism, and small molecule binding, paving the way for further drug development targeting the 12-LOX enzyme.

**Key Points:**

1. The first full-length structures of human arachidonate 12S-Lipoxygenase (12-LOX)
2. Reveals mechanisms of oligomeric and conformational states
3. Uncovers natural inhibitor of 12S-Lipoxygenase (12-lox)
4. Reveals a binding site of inhibitor ML355

## Introduction

Platelet activation is essential for maintaining haemostasis. However, uncontrolled platelet activation leads to abnormal clot formation and an increased risk of thrombosis and cardiovascular disease ^1,2^. Inhibition of platelet activation has been shown as an effective treatment that reduces the morbidity and mortality of cardiovascular ischemic events, such as myocardial infarction and stroke. Despite the use of antiplatelet therapies such as aspirin and P2Y_12_ receptor antagonists to reduce thrombotic risk, a high prevalence of ischemic events leading to unacceptable levels of morbidity and mortality remain. Due to this continued risk for thrombosis, the development of alternative therapies to further limit occlusive thrombotic events is warranted.

The enzyme, human 12-lipoxygenase (12-LOX, ALOX12), is highly expressed in platelets ^3^, and its activation leads to the production of 12-hydroperoxyeicosatetraenoic acid (12-HpETE), a prothrombotic oxylipin ^4,5^. The inhibition of 12-LOX prevents platelet activation ^6,7^, has a minimal effect on haemostasis, and does not promote increased bleeding, a common side effect of other antiplatelet therapies ^6,8,9^. The selective 12-LOX inhibitor, ML355 ^10^, has passed phase I clinical trials for the treatment of heparin-induced thrombocytopenia and has received fast-track designation by the FDA. Despite 12-LOX being a promising target for antiplatelet therapies there are no experimentally determined structures of the entire human 12-LOX; thus, limiting our understanding of the mechanism and regulation of 12-LOX activity. Although structures of other LOX isozymes have been determined by x-ray crystallography ^11–15^, they do not provide enough information to fully comprehend the mechanism of 12-LOX oligomerization and inhibitor binding.

Here we present the first high-resolution structures (1.7–2.8 Å) of human 12-LOX determined using cryo-electron microscopy (cryo-EM). We show that 12-LOX possesses a similar protein fold compared to other lipoxygenases^11–15^. From a single sample of 12-LOX, we were able to determine cryo-EM structures in multiple oligomeric forms from the monomer up to a hexamer. This observation is consistent with prior studies that demonstrated the existence of multiple oligomeric forms of lipoxygenases, including monomers, dimers and tetramers ^16–18^. Similar to human 15-LOX, we also captured 12-LOX in different conformational states that likely reflect the different parts of the catalytic cycle for this enzyme – the “open” and “closed” conformation ^11^. Due to the high-resolution features of the cryo-EM map, we were able to identify an endogenous 12-LOX inhibitor, a long-chain acyl-Coenzyme A, that co-purifies with the enzyme from mammalian cells. Finally, we were able to elucidate a putative allosteric binding site for the phase II inhibitor ML355. Collectively, we anticipate these structures guiding further research into the function of 12-LOX for platelet activation and promote further drug discovery efforts at this clinically relevant enzyme.

## Materials and Methods

### Expression and purification

The human 12-LOX with the N-terminal 6xHistidine tag, was cloned into pGen2 vector. The 1L of Expi293 cells grown in Expi293 expression media were transiently transfected using Polyethylenimine (PEI) “Max” (1mg of DNA and 3 ml of 1mg/ml PEI) and the expression was carried out in a shaking incubator for 5 days at 37°C, 7% CO2 in the presence of 1 *μ*M ML355. The cell pellet was thawed and resuspended in lysis buffer (20 mM HEPES pH 8.0, 30 mM NaCl, 0.5 mM TCEP, 1 mM MgCl2, 500 units benzonase, 0.2 mM PMSF, 5 μg/ml leupeptin, 5 μg/ml soybean trypsin inhibitor and 5 μM ML355) before being lysed by Dounce homogeniser using a tight pestle. Cell debris was removed by centrifugation (30,000 x g, 20 min, 4 °C) and the soluble portion was batch bound to Ni affinity resin for 1 hour at 4 °C. Ni affinity resin was packed into a glass column and washed with 10 column volumes of low salt wash buffer (20 mM HEPES pH 8.0, 150 mM NaCl, 0.5 mM TCEP, 10 mM Imidazole and 2 μM ML355) followed by 10 column volumes of high salt wash buffer (20 mM HEPES pH 8.0, 300 mM NaCl, 0.5 mM TCEP, 10 mM Imidazole and 2 μM ML355) before elution (20 mM HEPES pH 8.0, 150 mM NaCl, 0.5 mM TCEP, 500 mM imidazole and 5 μM ML355). Eluted protein was concentrated in an Amicon Ultra-15 50 KDa molecular mass cut-off centrifugal filter unit (Millipore, Burlington, MA, USA) and further purified by SEC on a Superdex 200 Increase 10/300 GL (Cytiva, Marlborough, MA, USA) in SEC buffer (20 mM HEPES pH 8.0, 150 mM NaCl, 0.5 mM TCEP and 5 μM ML355). Fractions were assessed for purity by SDS-PAGE, and fractions corresponding to tetramer or dimer were pooled separately; an additional 5 μM ML355 was added before concentrating, snap freezing in liquid nitrogen and storage at −80 °C.

### Grid preparation and cryo-EM imaging

Samples were thawed, diluted to 1 mg/ml, and incubated at 4 °C with an additional 40 μM ML355. UltrAufoil R1.2/1.3 300 mesh holey grids (Quantifoil GmbH, Großlöbichau, Germany) were glow discharged in air at 15 mA for 180 s using a Pelco EasyGlow. 3 μl of sample was applied to the grids at 4 °C and 100% humidity and plunge frozen in liquid ethane using a Vitrobot Mark IV (Thermo Fisher Scientific, USA) with a blot time of 3s and blot force of −3. Data were collected on a G1 Titan Krios microscope (Thermo Fisher Scientific, USA) equipped with S-FEG, a BioQuantum energy filter and K3 detector (Gatan, Pleasanton, California, USA). The Krios was operated at an accelerating voltage of 300 kV with a 50 μm C2 aperture, 100 μm objective aperture inserted, and zero-loss filtering with a slit width of 10 eV, at an indicated magnification of 105kX in nanoprobe EFTEM mode. Data were collected using aberration-free image shift (AFIS) with Thermo Fisher EPU software. Further parameters for each dataset can be found in **Table S1.**

### Tetramer cryo-EM image processing and model building

6896 movies collected at 0.82 Å/pix were motion corrected using UCSF MotionCor2 ^19^, and CTF parameters were estimated using GCTF^20^. 3.7 M particles were picked using autopicking within RELION v3.1^21,22^, and extracted particles were subject to multiple rounds of 2D and 3D classification with RELION v3.1. A set of particles (0.8 M) underwent 3D refinement, Bayesian polishing and CTF refinement before importing into cryoSPARC ^23^ where the last round of 2D classification and 3D heterogeneous refinement resulted in a final set of particles (0.5 M). The final set of particles underwent non-uniform 3D refinement with D2 symmetry, resulting in a 1.82 Å resolution map of the tetramer (FSC = 0.143, gold standard). Due to the high degree of flexibility between monomers, particles were subject to D2 symmetry expansion, and local refinement was performed on a monomer in cryoSPARC, which resulted in a 1.72 Å map. To model into the monomeric map, an initial predicted model was obtained from the AlphaFold Protein Structure Database^24,25^, rigid body fit into the density using UCSF ChimeraX^26^ and subject to repeated rounds of manual model building in COOT^27^ and real-space refinement in PHENIX^28^. The refined monomer model was used to model into the tetramer map similarly. Lastly, the models were quality assessed using MolProbity^29^ before PDB deposition.

### 3D variability analysis of tetramer

3D variability analysis (3DVA)^30^ was performed using cryoSPARC using the final symmetry expanded particles (1.8 M). 3DVA was initially performed masked on the entire tetramer with 3 principal components output to 20 frames and analysed in UCSF ChimeraX^26^. 3DVA was further performed masked on a monomer with 3 principal components output in clustered mode (5 clusters). A subset of particles (0.3 M) with clear density in the ligand binding site was subject to local refinements yielding a monomer map at 1.90 Å and a tetramer map at 2.05 Å. These maps were used to model oleoyl-CoA into the ligand binding site. Modelling was performed as above, oleoyl-CoA was obtained from the monomer library (3 letter code – 3VV), and geometry restraints were generated using the GRADE webserver^31^.

### Dimer cryo-EM image processing and model building

4812 movies at 0.82 Å/pix were motion corrected using UCSF MotionCor2^19^, and CTF parameters were estimated using GCTF^20^. 2.7 M particles were picked using Gautomatch using 2D projections of a previously processed 12-LOX dimer as a reference and extracted within RELION v3.1^21,22^ with 4x downsampling (80-pixel box size at 3.28 Å/pix). Particles were imported into cryoSPARC^23^ and subjected to 2D classification upon which the heterogeneity of the dataset became apparent. The 2D classes for the monomer, dimer, tetramer and the hexamer were manually selected and subjected to ab-initio reconstruction to create the starting models for each individual oligomeric state. The ab-initio models and an additional “junk” map were used for heterogeneous refinement of the entire dataset. From here on, the resulting particles for each oligomeric state were processed separately using similar strategies. First, each particle set was cleaned up using 2D classification. Then particle coordinates were imported back into Relion using pyem^32^ for re-extraction at full resolution (320-pixel box at 0.82 Å/pix). Each particle set was then subjected to 2D classification, heterogenous and non-uniform refinement in cryoSPARC followed by CTF refinement and Bayesian polishing in Relion v3.1 followed by final round of non-uniform refinement in cryoSPARC. Specific processing details for each of the oligomeric states are stated below. <u>The 12-LOX monomer</u> particle set contained 35,994 particles and yielded a 2.76 Å resolution map (FSC = 0.143) after a non-uniform 3D refinement. <u>The 12-LOX dimer</u> particle set contained 126,914 particles and yielded a 2.53 Å (FSC = 0.143) resolution map following a non-uniform 3D refinement. Each subunit of the dimer was further subjected to local refinement and 3D variability analysis to identify the least flexible classes. This yielded a “closed” (2.54 Å from 34,436 particles) and an “open” (2.33 Å from 126,914 particles) subunit conformations. <u>The 12-LOX tetramer</u> particle set contained 70,330 particles and yielded a 2.32 Å resolution map (FSC = 0.143) after a non-uniform 3D refinement in D2 symmetry. The local refinement of individual subunits following the symmetry expansion yielded a 2.1 Å map. <u>The 12-LOX hexamer</u> particle set contained 35,839 particles and yielded a 2.63 Å resolution map (FSC = 0.143) after a non-uniform 3D refinement in D3 symmetry. The local refinement of individual subunits following the symmetry expansion yielded a 2.24 Å map.

For modelling purposes of the 12-LOX dimer and the hexamer we created “composite” maps using Phenix suite^28^. To do this, for the 12-LOX dimer we combined the locally refined “open” and “closed” maps. For the 12-LOX hexamer we employed the locally refined single subunit map that was expanded using D3 symmetry. For modelling, the high-resolution model of the tetramer subunit (derived from the modelling into the highest resolution 1.72Å map of the tetramer) was rigid-body fit into the density maps of individual subunits of other oligomeric states using UCSF ChimeraX^26^ and subject to repeated rounds of manual model building in COOT^27^ and real-space refinement in PHENIX^28^. The ML355 and arachidonic acid (AA) were identified and manually built into 12-LOX. ML355 was generated from SMILES code, and geometry restraints were generated using the GRADE webserver^31^. AA was obtained from the monomer library (3 letter code – ACD). Lastly, the models were quality assessed using MolProbity^29^ before PDB deposition.

### Steady-State Kinetics

12-LOX reactions were performed at 22 °C in a 1 cm quartz cuvette containing 2 mL of 25 mM HEPES (pH 8.0) with AA varying from 0.25 to 25 μM. AA concentrations were determined by measuring the amount of oxylipins produced from complete reaction with soybean lipoxygenase-1 (SLO-1). Concentrations of 12-HpETE were determined by measuring the absorbance at 234 nm. Reactions were initiated by the addition ~20 μg of dimer 12-LOX and ~30 μg of tetramer 12-LOX and were monitored on a Perkin-Elmer Lambda 45 UV/Vis spectrophotometer. Product formation was determined by the increase in absorbance at 234 nm for 12-HpETE (ε_234nm_ = 25,000 M^-1^ cm^-1^). KaleidaGraph (Synergy) was used to fit initial rates (at less than 20% turnover), to the Michaelis-Menten equation for the calculation of kinetic parameters.

### IC_50_ Determination

The IC_50_ values for the 12-LOX specific inhibitor, ML355, against dimer 12-LOX and tetramer 12-LOX were determined in the same manner as the steady-state kinetic values. The reactions were carried out in 25 mM HEPES buffer (pH 8.0), 0.01% Triton X-100, and 10 μM AA. IC_50_ values were obtained by determining the enzymatic rate at nine inhibitor concentrations and plotting rate against their inhibitor concentration, followed by a hyperbolic saturation curve fit. The data used for the saturation curve fits were performed in duplicate or triplicate, depending on the quality of the data. Triton X-100 was used to ensure proper solubilization of the fatty acid and inhibitor. The acyl-CoAs were purchased from Avanti Polar Lipids. The IC_50_ values for the acyl-CoAs against dimer 12-LOX SEC peak were determined in a similar manner as described above.

### Synthetic Liposome Preparation and 12-LOX Binding

A lipid suspension was prepared from commercial sources with the following molar ratios: 99.9:0.1 DOPC: DSPE-PEG (i.e. DOPC). Each lipid mixture was dissolved in chloroform and the solutions were left under N2 for 20 min and placed in a vacuum chamber for at least 12 h at room temperature to remove all traces of solvent. Lipid mixtures were dissolved in 25 mM HEPES buffer (pH 8) to a concentration of 10 mg/mL and incubated in glass vials on a tube rotator for 1 h to facilitate homogenization. Liposomes were created using the literature protocol^33^ using a 100 nm filter. Liposome suspension volumes were adjusted to have a final concentration of 10 mg/mL and their size were determined by dynamic light scattering (DLS).

Liposome suspensions were placed on a DynaMag^™^-2 for 15 min at room temperature to bind the liposomes. Supernatant was removed and 1 mL of 25 mM HEPES buffer (pH 8) was added. This washing step was repeated five times, after which time 30 μg of dimer or tetramer were added to 1 mL of a 10 mg/mL liposome suspension. The sample was rocked over ice for 10 min and then placed on a DynaMag^™^-2 for 10 min. 25 μL of the supernatant was saved, and the beads were resuspended to a final volume of 1 mL. 25 μL of the resuspended beads were saved and both saved samples were subjected to SDS-PAGE and subsequent Western analysis. The antibodies were isolated from rabbit sera subjected to 12-LOX exposure (Pocono Rabbit Farm). Using ImageLab, ratios of the supernatant to pellet were determined. All conditions were done in triplicate.

## Results

### High-resolution structures of 12S-Lipoxygenase

The purification of 12-LOX from mammalian Expi293 cells yielded an oligomeric mixture of 12-LOX forms that were separated by size using size-exclusion chromatography (SEC) (**Fig. 1** and **Supp. Fig. 1**). The two main peak fractions corresponded to a dimer and a tetramer based on their size. Both SEC fractions displayed similar enzyme kinetics with a k_cat_ of 12 ± 0.9 s^-1^ for the dimer and 4.8 ± 0.2 s^-1^ for the tetramer. Both had a k_cat_/K_M_ value of 1.4 ± 0.1 s^-1^μM^-1^. The enzyme activities of both SEC fractions were inhibited by ML355 with IC_50_’s of 1.6 ± 0.3 *μ*M and 1.4 ± 0.3 *μ*M, for the dimer and tetramer, respectively (**Table S2**). Prior efforts to determine structures of 12-LOX using x-ray crystallography likely failed due to sample heterogeneity. Thus, we turned to cryo-EM to determine the structure of 12-LOX from each SEC fractions. To understand the binding mechanism of ML355 binding site, we added the inhibitor during protein expression and purification.

**Figure 1.**
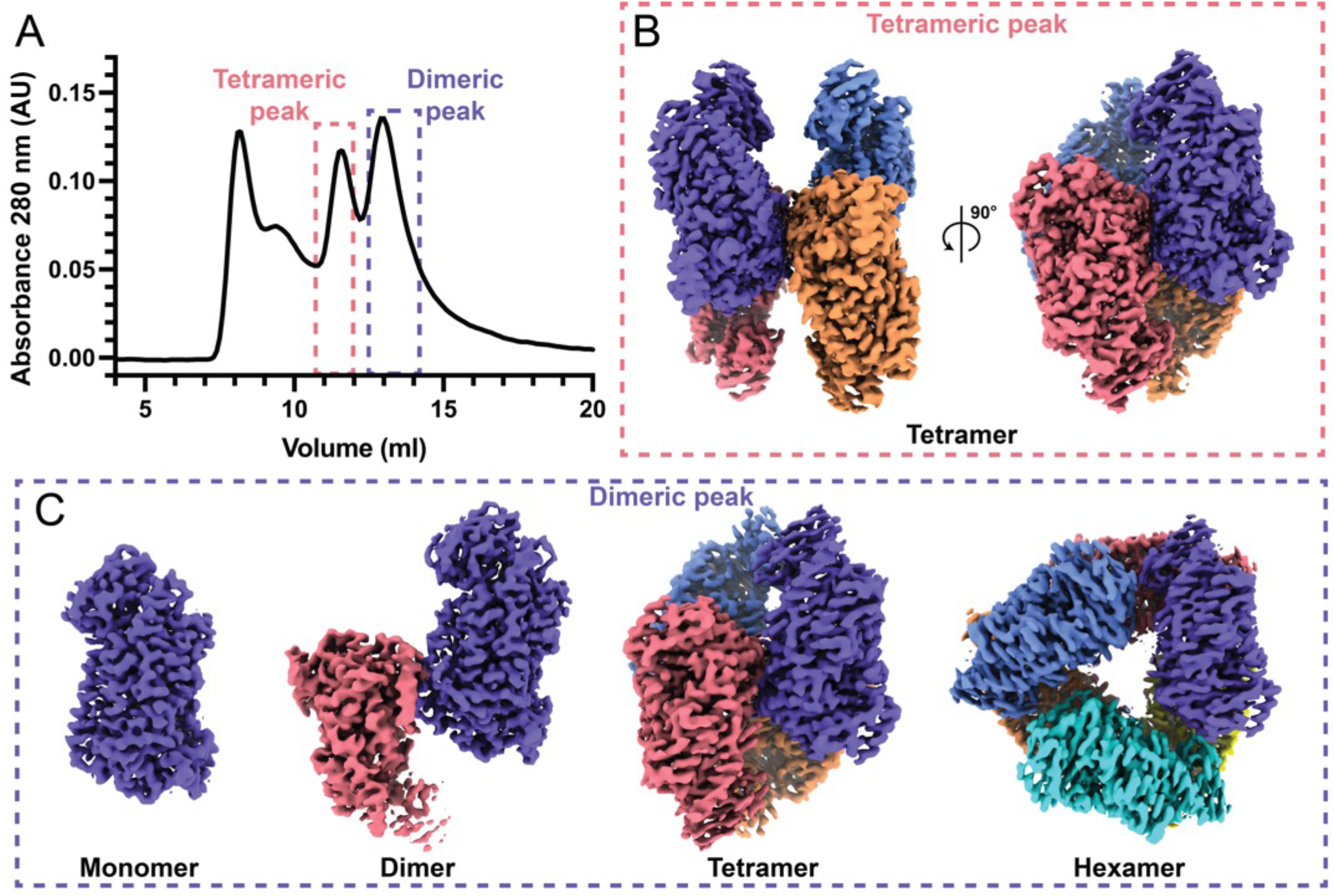
Different oligomeric forms of 12-LOX. (A) Size exclusion chromatography (SEC) UV absorbance trace (280 nM absorbance) from 12-LOX purification. 12-LOX separated as two distinct peaks, one corresponding to a tetramer (red box) and the other a dimer (blue box). (B) Cryo-EM map of a 12-LOX tetramer from “tetrameric” SEC peak. (C) Cryo-EM maps of different 12-LOX oligomers resolved from “dimeric” SEC peak.

Unexpectedly, the dimer peak from the SEC 12-LOX purification gave rise to multiple high-resolution cryo-EM structures of 12-LOX in different oligomeric forms including monomers (2.8 Å), dimers (2.5 Å), tetramers (2.3 Å), and hexamers (2.6 Å), all from the same imaged grid (**Fig. 1**, **Supp. Fig. 2, 4** and **Table S1**). In contrast, the tetramer peak yielded a structure of only the 12-LOX tetramer (**Fig. 1**, **Supp. Fig. 3, 4** and **Table S1**) with an overall resolution of 1.8 Å. All oligomeric forms of 12-LOX exhibited significant intermolecular flexibility determined by 3D variability analysis (3DVA) in cryoSPARC (**Supp. Movies 1-3)^30^**. Thus, we employed local refinement (cryoSPARC) to improve the map resolution and quality for the individual subunits (**Supp. Fig. 2-4**) within each oligomer. For the tetramer peak sample, this improved the resolution to 1.7 Å allowing for accurate model building of the full-length 12-LOX (residues G2 to I663) (**Fig. 2A-B**).

**Figure 2.**
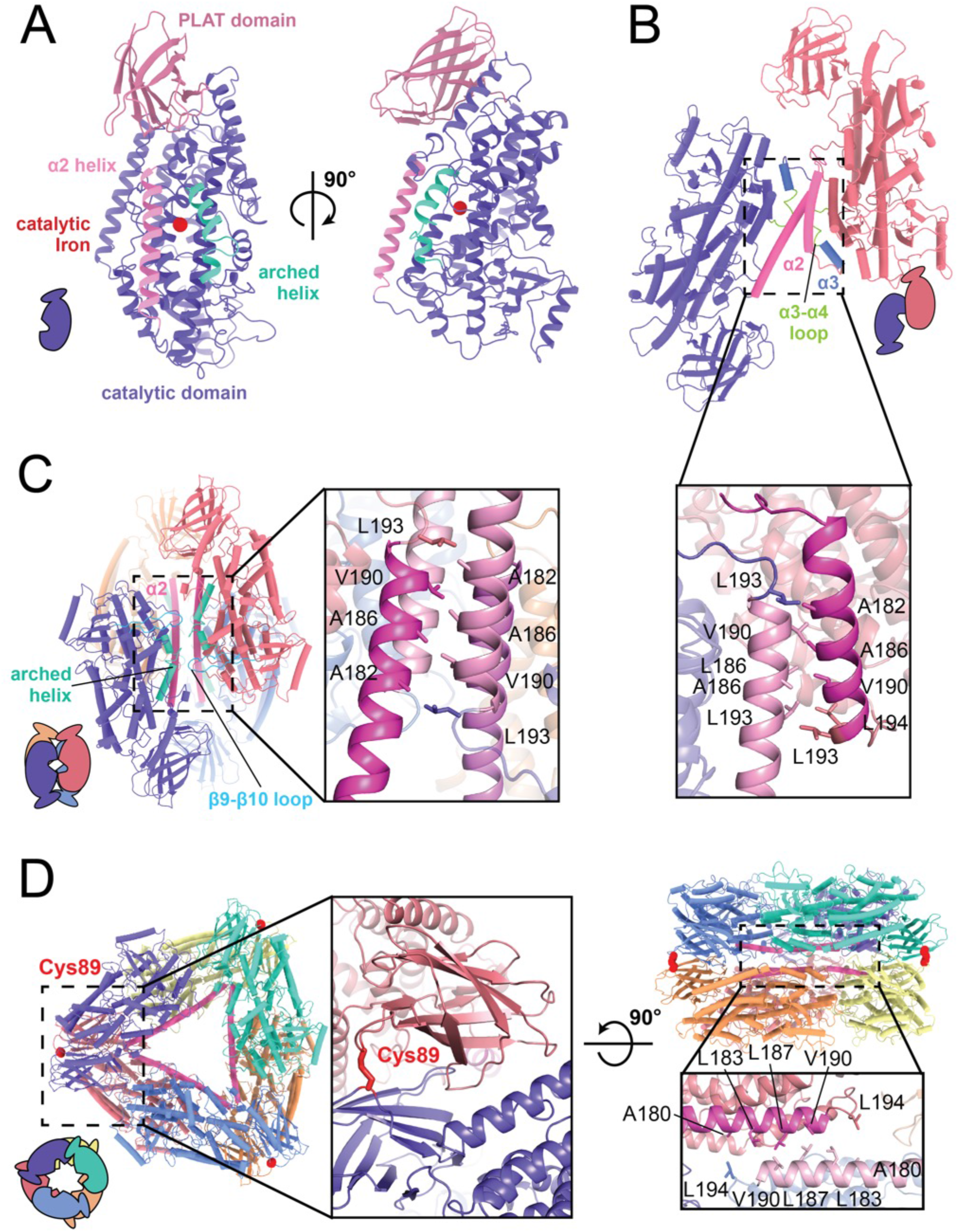
Oligomeric structures of 12-LOX. Models of 12-LOX as (A) monomer, (B) dimer, (C) Tetramer and (D) hexamer. Each subunit represented in a different colour and the α2-helix coloured in pink. Graphical representation of each oligomeric state in bottom left and insets details the oligomeric interface. Interacting amino acids are shown as sticks. (D) Cys39 (in red) contributes a disulphide bridge to the interface of the hexamer. Fe atom is shown as a red sphere.

The structural architecture of 12-LOX is typical of lipoxygenases with the N-terminal β-barrel PLAT (polycystin-1-lipoxygenase α-toxin) domain and a C-terminal α-helical catalytic domain (**Fig. 2A**). Structural alignments with other LOX structures (**Supp. Fig. 5A**) revealed markedly similar folds with root mean square deviations (RMSDs) <1Å with the largest variation occurring in the PLAT domain (**Supp. Fig. 5B**). The active site of 12-LOX is located in the catalytic domain, where a catalytic iron atom is coordinated by three conserved histidine residues (H360, H365 and H540) as well as N544 and the carboxyl C-terminus of I663 (**Supp. Fig. 5C-D**). Next to the catalytic iron is the typical LOX U-shaped lipid-binding pocket that is lined with hydrophobic residues. The entrance to the active site is bordered by an arched helix and an α2-helix in an extended conformation (**Fig. 2A** and **Supp. Fig. 5C-D**).

### Oligomeric states of 12S-lipoxygenase

The biological unit of the 12-LOX oligomers appears to be a dimer arranged “head-to-toe”. The tetramer and hexamer are made of a dimer of dimers and a trimer of dimers, respectively (**Fig. 1** and **2**). The overall dimer substructure in each oligomeric form is maintained mainly through Van der Waals interactions between the α2-α4 helixes and the α3-α4 loop (**Fig. 2B-D**). The dimer substructures of the 12-LOX tetramer and hexamer were virtually identical (RMSDs 0.78Å), while the individual subunits of the dimer were rotated by 30°, due to change in the conformation of the α2-helix (described below) (**Supp.Fig. 6**). Higher-level oligomerisation in 12-LOX tetramers was maintained through additional Van der Waals interactions of the α2-helix and hydrogen bond interactions of the arched helix and β9-β10 loop between neighbouring subunits (**Fig. 2C**). The architecture of the hexamers was supported by an additional hydrogen bonding network and a disulphide bond (C89-C89) between neighbouring PLAT domains (**Fig. 2D).**

The oligomerisation of 12-LOX affects the accessibility of the active site to bulk solvent. The entrance to the 12-LOX catalytic site is defined by the α2- and arched helixes and is in the same plane as the predicted membrane-binding residues W70/L71/A180^34^. The entrance is accessible to solvent in the 12-LOX monomer, dimer, and hexamer, but is obscured when the two dimers associate to form a tetramer. (**Supp. Fig. 7**).

To investigate whether the oligomerisation affects 12-LOX membrane binding we tested the dimer and the tetramer SEC fractions (**Fig. 1**) for their ability to bind artificial DOPC liposomes. Both 12-LOX preparations bind liposomes with similar extent (21+/-6% and 36+/-6% for dimers and tetramer peak, respectively) (**Supp Fig. 1C**).

### Conformational changes of 12S-Lipoxygenase

The structure of the 12-LOX monomer in all oligomeric forms were similar, except for the 12-LOX dimer. Similar to the arrangement within the 12-LOX tetramer and the hexamer, the monomers in the dimer are arranged “head-to-toe”, with most of the contacts mediated via hydrophobic interactions between the α2-helixes (**Fig. 2B**), previously determined by HDX-MS^35^. However, contrary to the protein chains in the tetramer and hexamer, the individual subunits in a dimer are not equivalent. Instead, they adopt either an “open” conformation (as observed in the 12-LOX monomer, tetramer, and a hexamer) or a “closed” conformation, predominantly facilitated by a large-scale motion of the α2-helix and corresponding rearrangements of the neighbouring loops (**Fig. 3**). In the open conformation, the α2-helix forms a long single helix that lies at the edge of the active site. Conversely, in the closed conformation, the α2-helix undergoes a rigid 23° pivot and rotation that blocks the entrance to the active site reducing its internal volume (**Fig. 3B**). The conformational change of the α2-helix also leads to a 30° rotation of the two monomers relative to each other and relative to the dimer substructure observed in the 12-LOX tetramers and hexamers (**Supp. Fig. 6**).

**Figure 3.**
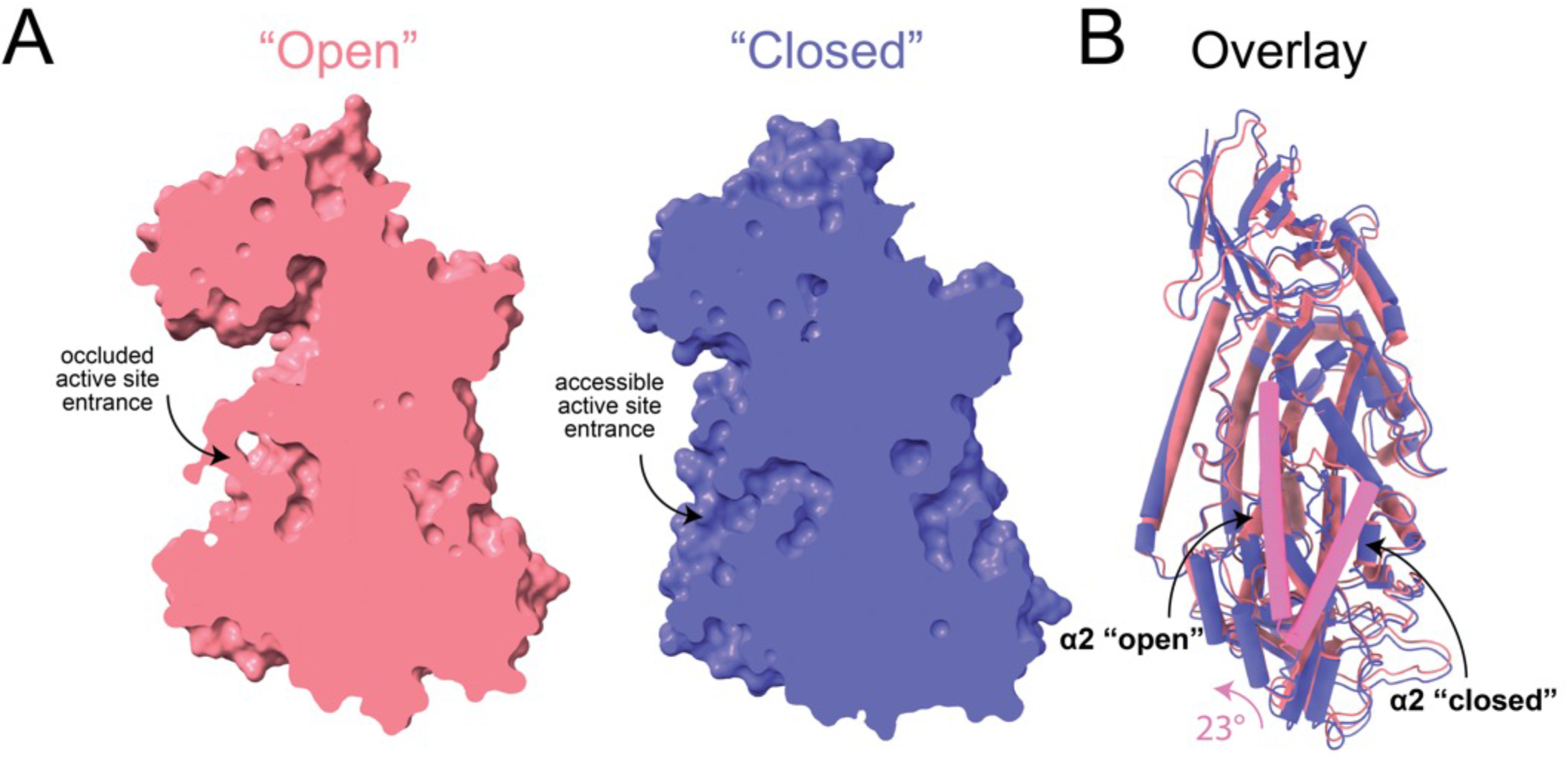
Conformational changes in the 12-LOX dimer. (A) Surface representation of 12-LOX in the “closed” (left) and “open” states (right) showing the active site cavity of the dimer 12-LOX subunits. In the open conformation this cavity is occupied by a small molecule. (B) An alignment of “open” and “closed” states shows a 23°C rotation and unwinding of the N-terminal residues of the α2-helix. The structures are shown as cartoons with cylindrical helixes, the α2-helix is in pink.

### Natural inhibition of 12S-Lipoxygenase by long chain fatty acid acyl-CoAs

In all of our 12-LOX structures the active site of the 12-LOX subunits in the open conformation was occupied by extra density in the cryo-EM maps (**Supp. Fig. 8**), suggesting the presence of a bound ligand. The shape of the density varied between oligomeric forms suggesting different ligands. The high-resolution of the tetramer (1.7 Å) and the hexamer (2.3 Å) cryo-EM maps allowed us to model ligands into these densities with high confidence. However, due to the lower resolution of the monomer and dimer cryo-EM maps, we were not able to confidently identify the bound molecule. We hypothesized the observed densities were either ML355 or an endogenous lipid(s) co-purified from Expi293 cells.

The 12-LOX tetramer is made of a dimer of dimers. The subunits at the inter-dimer interface face each other with their lipid binding sites (**Fig. 4A**). Within each of the U-shaped pockets, we observed a density that resembled a lipid tail. The lipid density extended out of the binding site, spanning the gap between two neighbouring subunits (**Supp. Fig. 9A**). This density was also present in the apo 12-LOX tetramer samples that were expressed and purified in the absence of ML355 suggesting the ligand was co-purified from the HEK293 cells (data not shown). To improve the resolution of the cryo-EM maps further, we performed a 3D variability analysis (3DVA) on individual subunits within a tetramer (**Sup Fig. 9B, Supp. Movie 2).** Using the cluster mode of the 3DVA, we were able to separate the protein chains that were fully occupied with the molecule and reconstruct the corresponding 12-LOX subunits and a full tetramer to a resolution of 1.9 Å and 2.05 Å, respectively (**Fig. 4A-C**). Furthermore, the 3DVA revealed that the lipid is only bound to one of the subunits at the inter-dimer interface at a time (thus averaging to ½ lipid occupancy in the entire 12-LOX tetramer) (**Supp. Fig. 9C, Supp. Movie 4**). In contrast, the opposite subunit was mostly empty with some weak non-continuous density in the active site that could represent another unidentified lipid or incomplete separation of the occupied vs. unoccupied subunits during 3DVA.

**Figure 4.**
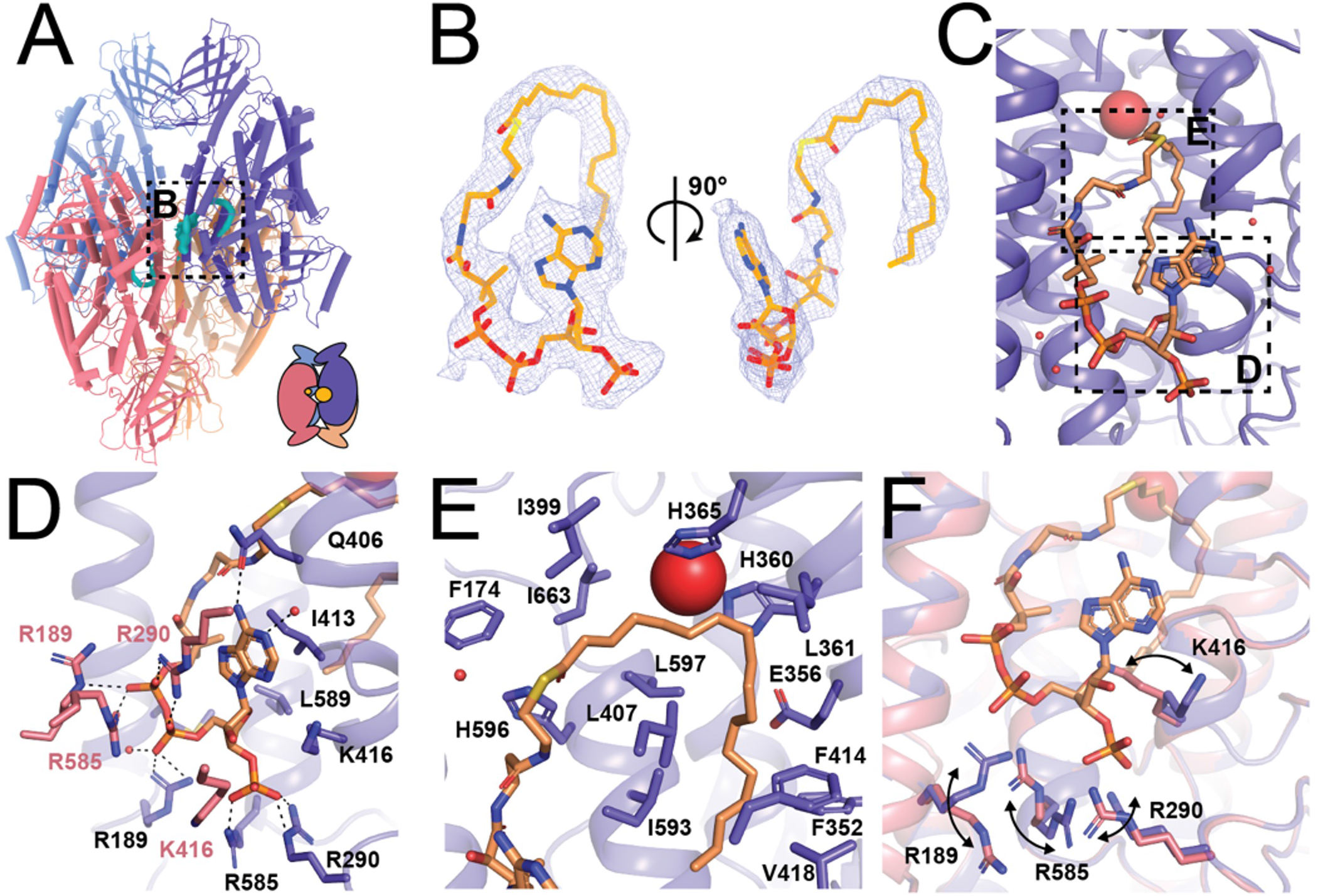
Acetyl-CoA binding site in the 12-LOX tetramer. (A) Model of tetrameric 12-LOX, with density in the catalytic site shown as cyan volume. Graphical representation is in right corner. (B) Acyl-CoA model and the density. Density is shown as wire mesh, the model is in sticks coloured by heteroatoms. (C) Model of acetyl-CoA within the catalytic site of 12-LOX. (D-E) 12-LOX residues that contact the acetyl-CoA (orange) shown as sticks. (D) Contacts of the adenosine tri-phosphate group. (E) Contacts of the acetyl tail. (F) Conformational change of residues in contact with the acetyl-CoA. Acetyl-CoA bound subunit in purple and unbound in pink. Fe atom is shown as a red sphere.

Because of the high resolution and quality of the density map, we were able to identify the lipid as a fatty acid acyl-CoA ester with a tail approximately 18 carbons long and unknown saturation (oleoyl-CoA was used for modelling purposes) (**Fig. 4B**). The CoA headgroup is positioned at the inter-dimer interface at the entrance to the catalytic site, between the α2-helix and the arched helix with the fatty acid tail extending into the U-shaped hydrophobic cavity (**Fig. 4C**). The purine group of CoA forms CH-pi interactions with I413, a hydrogen bond with Q406 of the arched helix, and cation-pi interactions with the neighbouring molecule’s R290 (**Fig. 4D**). The three phosphate groups form electrostatic interactions with R189, R290, and R585 of the bound 12-LOX, as well as R189, R290, K416 and R585 of the neighbouring 12-LOX subunit. Overlay of these two subunits reveals that the polar residues at the dimeric interface undergo significant rearrangement to better accommodate the interaction with oleoyl-CoA (**Fig. 4F).** The fatty acid tail extends into the catalytic site, forming extensive hydrophobic contacts (**Fig. 4E**).

Due to the chemical lability of the acyl-CoA’s thioester and hence difficulty in detection by mass spectrometry, we set out to confirm our structural findings by determining whether fatty acid acyl-CoAs inhibit 12-LOX. We tested a panel of long chain acyl-CoAs with different lipid tail length and saturation to determine their ability to inhibit 12-LOX catalysis (**Table S3**). The data demonstrate that 12-LOX inhibition by acyl-CoAs is dependent on both their length and saturation status, with oleoyl-CoA (18:1) being the most potent inhibitor with an IC_50_ of 32 ± 4 *μ*M. Acyl-CoAs with similar carbon length (C18) but different levels of unsaturation, stearoyl-CoA (18:0) and gamma-linoleoyl-CoA (18:3), did not inhibit 12-LOX. arachidonoyl-CoA (20:4) weakly inhibits 12-LOX, with an IC_50_ of 110 ± 20 *μ*M, but shorter acyl-CoAs, such as palmitoyl-CoA (16:0) and palmitoleoyl-CoA (16:1), did not inhibit 12-LOX. None of the tested acyl-CoAs were substrates for 12-LOX. These data confirm that oleyl-CoA is the most potent inhibitor of 12-LOX, although the exact nature of the bound acyl-CoA in the structure is unconfirmed.

### ML355 binding of 12-LOX

In contrast to the tetramer structure, cryo-EM density within the active site of the 12-LOX hexamer was identical across subunits and was distinct from the acyl-CoA. Moreover, the density was of two independent molecules that could be perfectly fit with AA and ML355 (**Fig. 5A-C**). The AA molecule occupies the U-shaped hydrophobic cavity that was occupied by the fatty acid tail of the acyl-CoA in the 12-LOX tetramer. The carboxyl group of AA interacts with 12-LOX via a H-bond with H596, as predicted ^36^, positioning the C11-C12 double bond in the vicinity of the catalytic iron. The remainder of the contacts are from Van der Waals interactions with hydrophobic residues lining the channel of the active site (**Fig. 5E**). The position of AA is nearly identical to that of the anaerobic structure of coral 8R-LOX ^37^.

**Figure 5.**
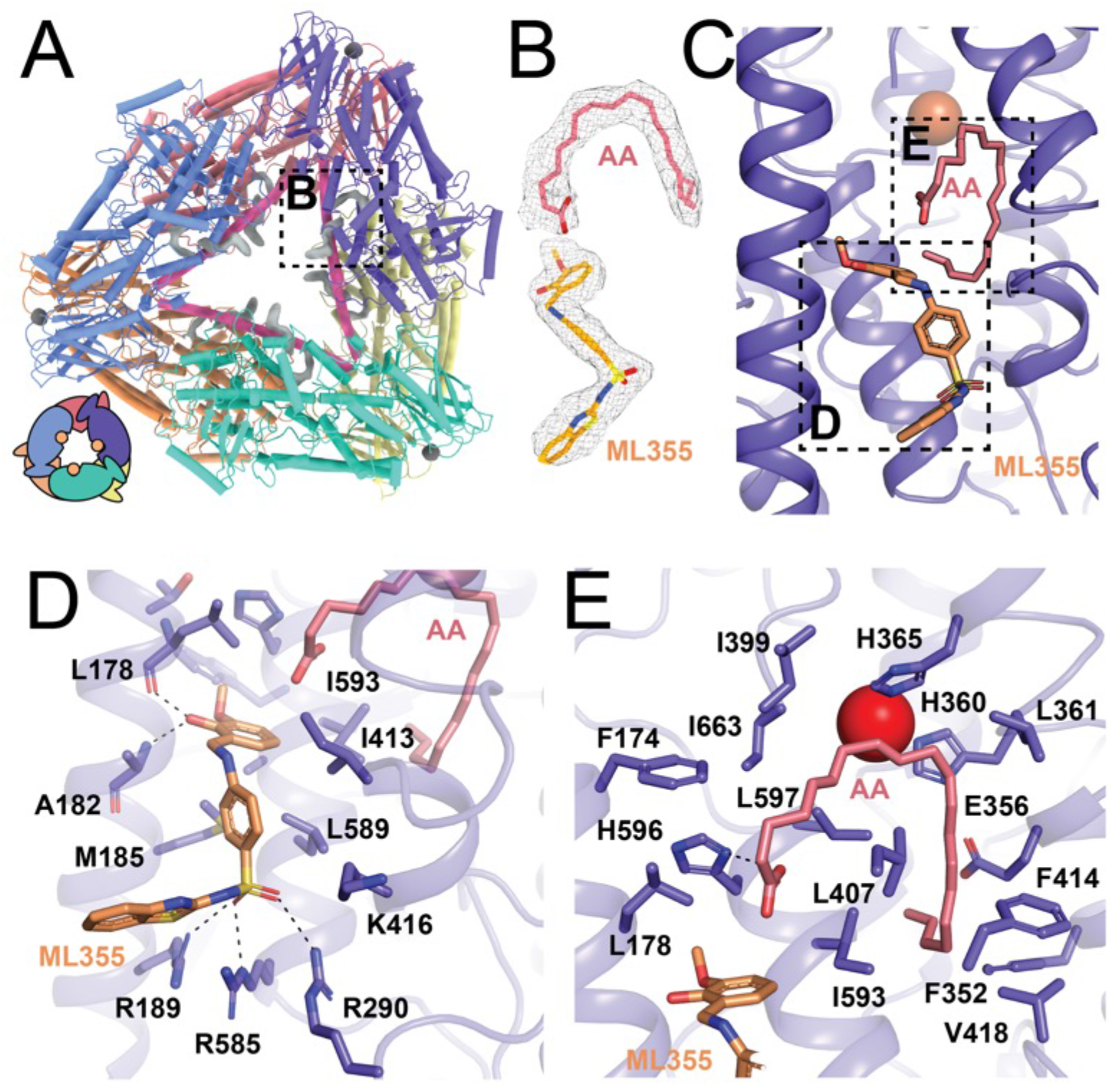
ML355 and arachidonic acid (AA) binding sites in the 12-LOX hexamer. (A) Model of hexameric 12-LOX with density in the catalytic site shown as grey volume. Graphical representation in the bottom left. (B) Density for AA and ML355. Density is shown as wire mesh, the models is in sticks (orange for ML355 and pink for AA) coloured by heteroatoms. (C) Model of 12-LOX bound to ML355 and AA. (D-E) 12-LOX residues that contact (D) ML355 (orange) and (E) AA (pink) shown as sticks. Fe atom is shown as a red sphere.

Docking and mutagenesis studies predicted ML355 to bind deep in the 12-LOX active site ^38^, but our cryo-EM density maps showed no evidence of ML355 occupying that region.

Unexpectedly, however, a molecule of ML355 perfectly fit into the EM density found at the entrance to the active site in the hexamer. Interactions of ML355 with 12-LOX include the hydroxyl group of the 2-hydroxy-3-methoxyphenyl moiety forming H-bonds with the backbone carbonyl of L178 and amide of A182 (**Fig. 5D**). The sulphur of benzothiazole ring forms a H-bond with R189 with the sulphonyl group within H-bonding distance to R290 and R585. The sulphonyl interactions of ML355 mimic the interactions observed with the phosphates from oleoyl-CoA and residues R189, R290, and R585. The rest of the molecule forms Van der Waals interactions with M185, I413, L589 and I 5993 (**Fig. 5D).**

## Discussion

To our knowledge, this is the first study to use cryo-EM to determine high-resolution structures of lipoxygenases. Compared to x-ray crystallography, the ability of cryo-EM to separate heterogeneous samples into discrete populations allowed us to gain unique insights into the nature of different 12-LOX oligomeric states. Human LOXs show diversity in their oligomeric state. Both 5-LOX and 15-LOX function as monomers but dimerize at higher protein and salt concentrations^17^. On the contrary, 12-LOX is primarily dimeric in solution^39^, but can also form larger aggregates^16^. Other studies suggested that most human LOXs can form high-order oligomers in solution^40^. Our structures provide the first high-resolution insights into the diversity of 12-LOX oligomeric forms that can likely be extended to other LOXs.

SAXS experiments predicted that all LOX dimers (12-LOX^16^, 15-LOX^18^ and 5-LOX^17^) have a similar organisation, including the “head-to-toe” arrangement of individual monomers that are interacting through their α2-helixes. Prior to our structures, such an arrangement was only observed in x-ray structures of rabbit 15-LOX-1^11,13^ and human 15-LOX-2^41^. While the overall dimer organisation was similar between all 3 enzymes (“head-to-toe” orientation and α2 helix-mediated interactions), the relative position of individual subunits varies, owing to differences in specific interacting residues.

While there is biochemical evidence for LOX oligomerisation ^16,18^, prior to this study, there was no structural information on LOX oligomers beyond dimers. Cryo-EM allowed us to determine the structures of 12-LOX tetramers and hexamers. Interestingly, both are made from the dimer building blocks that further oligomerise either into the dimer of dimers or trimer of dimers. Prior studies suggested that reducing agents might prevent the oligomerisation of 12-LOX ^16^, proposing that higher-order oligomers form through intramolecular disulphide bonds upon protein oxidation. We could only confirm this observation with the 12-LOX hexamer, observing a single disulphide bond (C89-C89). Other interactions in both the tetramer and a hexamer included an extensive network of hydrophobic interactions and hydrogen bonds, respectively. As such, the assembly of 12-LOX into monomers, dimers, and tetramers is independent of the oxidation state of the enzyme, while higher-molecular oligomers could represent a change to the oxidative environment of the cell.

The oligomerisation of 12-LOX might be a regulatory mechanism for enzyme activity and/or membrane binding as the accessibility of the active site varies between different oligomeric forms. The predicted membrane-binding residues for the 12-LOX are located in the same plane as the entrance to the 12-LOX binding site. Interestingly, the membrane binding surface within the 12-LOX dimer building block (present in dimers, tetramers and hexamers) is located within the same surface plane. However, in the tetramer the membrane-binding surface and active site entrance are further sequestered by interdimer contacts. As such, they might represent inactive states or storage pools for the enzyme. While we did not observe significant differences in AA oxidation rates or DOPC liposome binding between the dimer and tetramer SEC peaks used for cryo-EM data collection, the data could possibly be explained by our preparations being extremely heterogeneous and containing a mixture of the 12-LOX oligomeric forms (based on the structural data). Thus, further analysis using isolated 12-LOX oligomeric forms is necessary to better understand their role in membrane binding and catalysis.

Intriguingly, the higher-order oligomers of 15-LOX-1 were found to induce pore formation in the lead to organelle clearance during erythrocyte maturation^42^. The two-ring arrangement of the 12-LOX hexamer creates a channel with a diameter of ~30 Å. While the physiological role of this oligomeric species of 12-LOX requires further investigation, it is tempting to speculate that similar conformations might exist in other LOXs.

The protein chains in the 12-LOX monomer, tetramer, and hexamer adopt “open” conformation characterised by an extended a2-helix that pack along the entrance to the active site. Such an α2 conformation is seen in many of the LOX structures, including coral 8R-LOX ^37^, human 15-LOX-2 ^14^ and porcine 12-LOX (ALOX15)^15^. In the 12-LOX dimer, one subunit adopts an “open” conformation while another undergoes significant conformational change involving a large-scale α2 movement. The alterations to the extended α2 conformation were observed previously in crystal structures of stable 5-LOX ^12,43–45^ (broken or disordered α2) and 15-LOX-1 ^11,13^ (large-scale α2 movement). The 15-LOX-1 and now the 12-LOX are the only LOXs that were captured forming non-symmetrical dimers with one subunit in the “open” and one in the “closed” conformations. While the conformational change leading to the formation of the “closed” conformation differs in the degree of the α2 movement and the subunit rotation relative to each other, both result in the closure of the entrance to the active site.

The conformational change between the “open” and “closed” subunits in LOX dimers might be linked to its catalytic cycle. Thus, only the “closed” structure of the 15-LOX-1 was bound to an inhibitor (3-(2-octylphenyl)-propanoic acid), while the “open” subunit was empty^11^. Vice versa, the molecular dynamics simulation of a 15-LOX-1 dimer bound to a single molecule of AA showed the closure of the α2 over the active site entrance^18^. Similar to the 15-LOX-1 structure, our structures of 12-LOX oligomers demonstrate half occupancy of their active sites. In the 12-LOX dimer, only the active site of the “open” subunit is occupied by what appears to be a lipid density. Likewise, only one subunit of the dimer substructures of the tetramers is bound to the acyl-CoA. This suggests that only half of the oligomeric subunits may be active at any given time, while the other subunit serves a regulatory role. This mechanism could be responsible for differences in inhibitor binding observed previously between the dimeric and monomeric 12-LOX (converted by introducing L183E/L187E mutations). Only the dimer showed inhibition by ML355 (Ki = 0.43 *μ*M), while monomeric 12-LOX was unaffected^35^. Interestingly, similar behaviour has been previously described for another class of lipid-modifying enzymes, cyclooxygenases (COX). COX forms homodimers where one monomer acts as an ‘‘allosteric’’ subunit modulating the catalytic efficiency of its partner monomer, leading to ½ site occupancy during catalysis^46,47^.

One of the unexpected findings was the presence of the fatty acid acyl-CoA molecule in the 12-LOX tetramer that co-purified with our enzyme from Expi293 cells. Fatty acid acyl-CoA derivatives have long been known to inhibit platelet aggregation^48,49^ in a chain, length, and saturation-dependent manner. Specifically, the medium-chain acyl-CoA (palmitoyl, stearoyl, oleoyl and linoleoyl) inhibit lipoxygenase activity in platelets at concentrations ranging from 10 to 50 *μ*M^50^. We have confirmed that these acyl-CoAs directly inhibit 12-LOX at micromolar concentrations. Considering that the levels of acyl-CoAs within the cell could reach micromolar concentration ^51^, the long chain acyl-CoAs could be physiologically important regulators of 12-LOX function in the cell. The effect of acyl-CoA on platelet aggregation is thought to be mediated through P2Y1 and P2Y12 receptors ^52^. However, with the discovery that fatty acid acyl-CoA directly binds and inhibits 12-LOX, it might be possible that the inhibition of 12-LOX could also contribute to this process.

Despite the presence of ML355 during the expression and purification of 12-LOX, ML355 was only bound in the hexameric form of 12-LOX. It is likely, that ML355 was competed out in the other oligomeric forms due to the presence of endogenous lipids. The observed pose of ML355 is in contradiction to previously published docking/ mutagenesis studies that predicted ML355 binding deep in the 12-LOX active site ^38^. However, the simultaneous binding of ML355 and AA observed in our structure could explain the “mixed” mode of ML355 inhibition described previously^53^. Nevertheless, future studies are needed delineate the mechanism of ML355 inhibition with respect to different oligomeric forms of the enzyme along with the role of endogenous inhibitors that may or may not be present in platelets.

In conclusion, this study presents the first high-resolution cryo-EM structures of 12-LOX in multiple oligomeric forms, provides the first structural information on the clinically relevant 12-LOX inhibitor ML355, shows evidence for conformational changes that might accompany the 12-LOX catalytic cycle, and demonstrates that acyl-CoA can serve as endogenous 12-LOX inhibitor. This structural information will aid future studies of 12-LOX biology and its contribution to platelet activation and facilitate structure-based drug discovery efforts on a therapeutically validated enzyme.

## Acknowledgements

The work was supported by the funding from WEHI, The University of Melbourne and the estate of Akos and Marjorie Talon. A.G is a CSL Centenary Fellow. DMT is a National Health and Medical Research Council of Australia (NHMRC) Early Career Investigator fellow. M.H is supported by the National Institute of Health (NIH) grant R35 GM131835. We acknowledge use of facilities within the Monash Ramaciotti Cryo-EM platform and Ian Holmes Imaging Centre at the Bio21 Molecular Science and Biotechnology Institute. The computational work was supported by the MASSIVE HPC facility (https://www.massive.org.au).

## Author contributions

A.G. developed protein purification strategy, performed protein expression and negative stain transmission EM. A.G. and J.I.M purified the protein. H.V. vitrified the sample and performed image acquisition within the Monash EM facility. J.I.M., K.A.B and A.G. performed cryo-EM data processing, model building, refinement and validation. M.T. performed enzyme kinetics, inhibition and liposome binding. M.H., T.D., participated in experimental design and result interpretation. A.G., D.M.T, J.I.M. and K.A.B wrote the manuscript with contributions from all authors. A.G. and D.M.T. supervised the project.

## Data availability

Atomic coordinates and the cryo-EM density maps have been deposited in the Protein Data Bank and the Electron Microscopy Data Bank. The accession codes are 8GHB and EMD-40039 for 12-LOX monomers; 8GHC and EMD-40040 for dimers; 8GHE and EMD-40042 for tetramers and 8GHD and EMD-40041 for hexamers.

