## Supplemental Figures and Tables for "Cryo-EM structures of human arachidonate 12S-Lipoxygenase (12-LOX) bound to endogenous and exogenous inhibitors"

#### Supplementary Figure 1

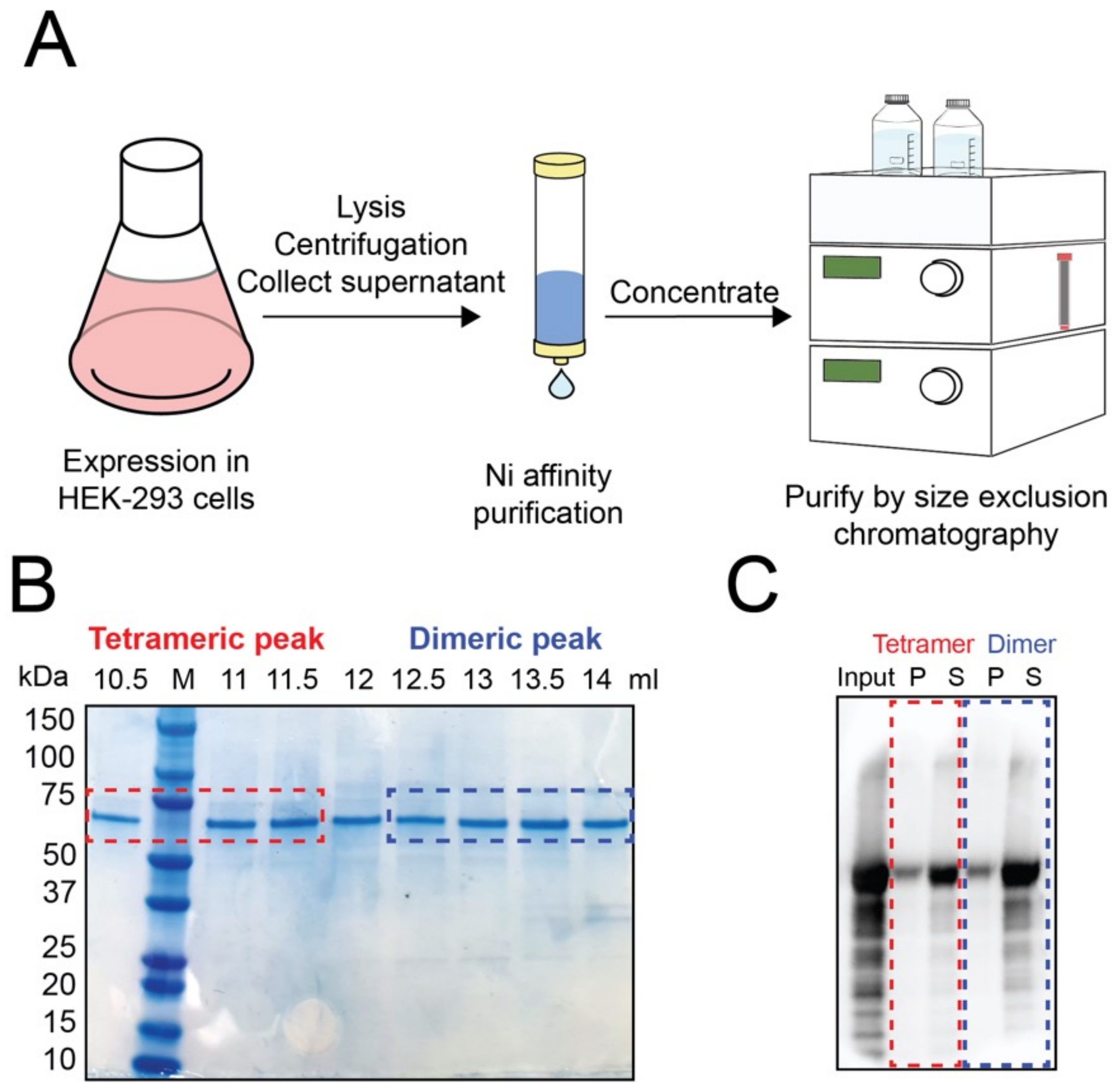


**Supplementary Figure 1. Expression and purification of 12S-Lipoxygenase (12-LOX).** (A) Graphical representation of 12-LOX expression and purification. 12-LOX was expressed in HEK-293 cells and purified by Ni-affinity and size exclusion chromatography (SEC). (B) SDS-PAGE of fractions following SEC. (C) Western Blot from a representative liposome binding experiment. P- pellet fraction; S-soluble fraction.

#### Supplementary Figure 2

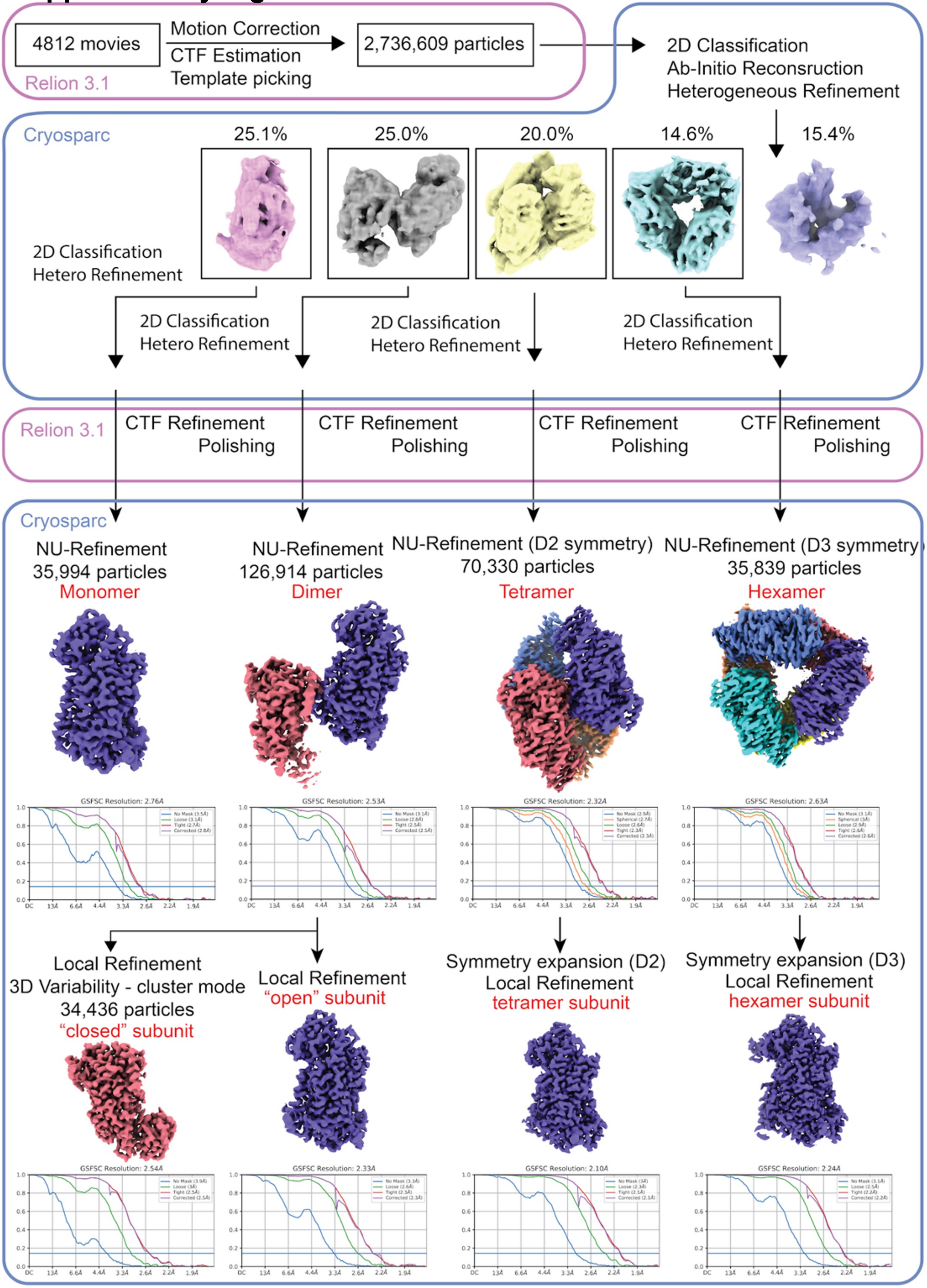


**Supplementary Figure 2. Cryo-EM data processing for dimer 12-LOX.** Workflow to achieve high-resolution structures of dimeric 12-LOX.

#### Supplementary Figure 3

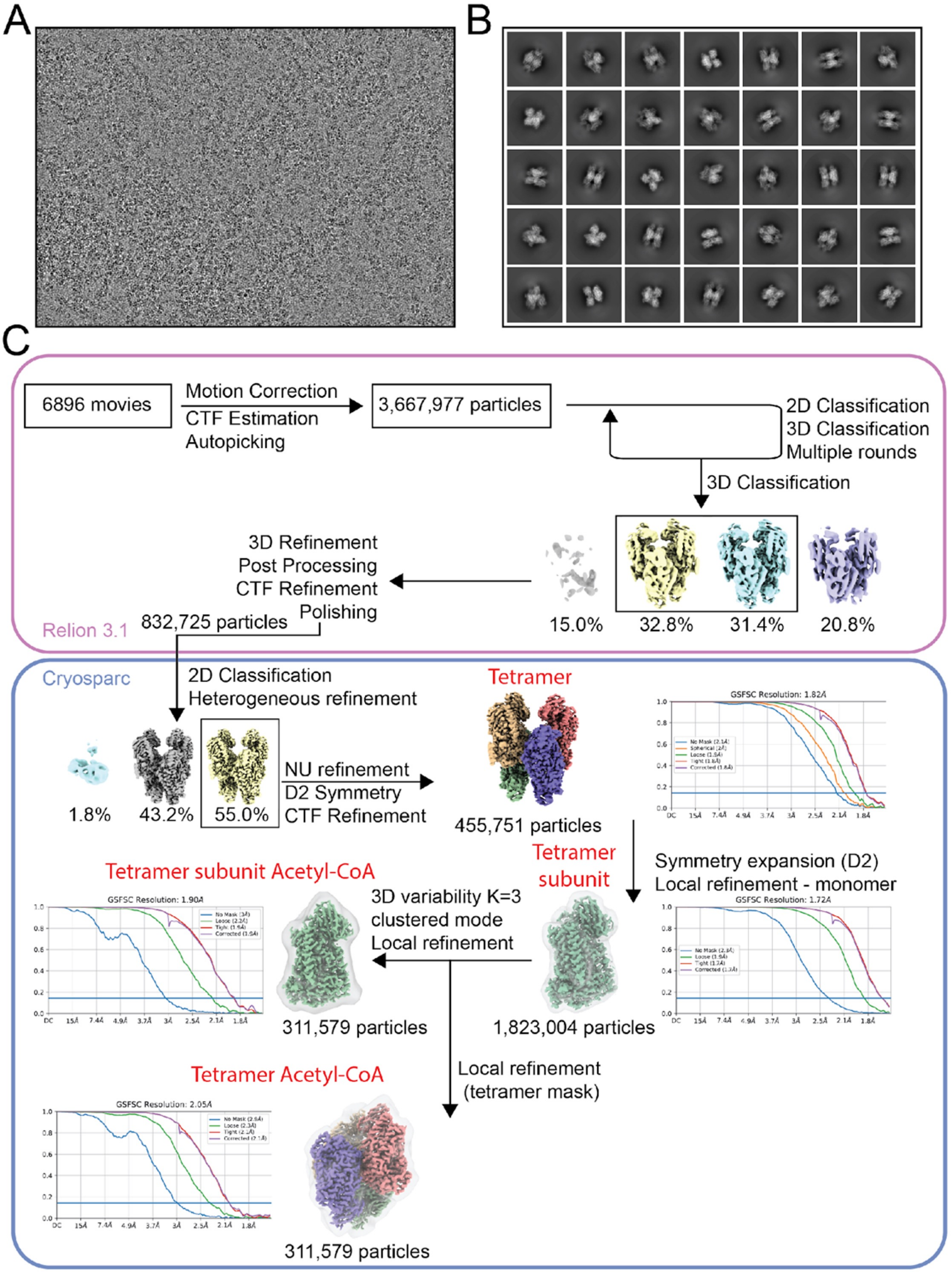


**Supplementary Figure 3. Cryo-EM data processing for tetramer 12-LOX.** (A) Representative micrograph and (B) representative 2D classifications. (C) Workflow to achieve high-resolution structures of tetrameric

### Supplementary Figure 4

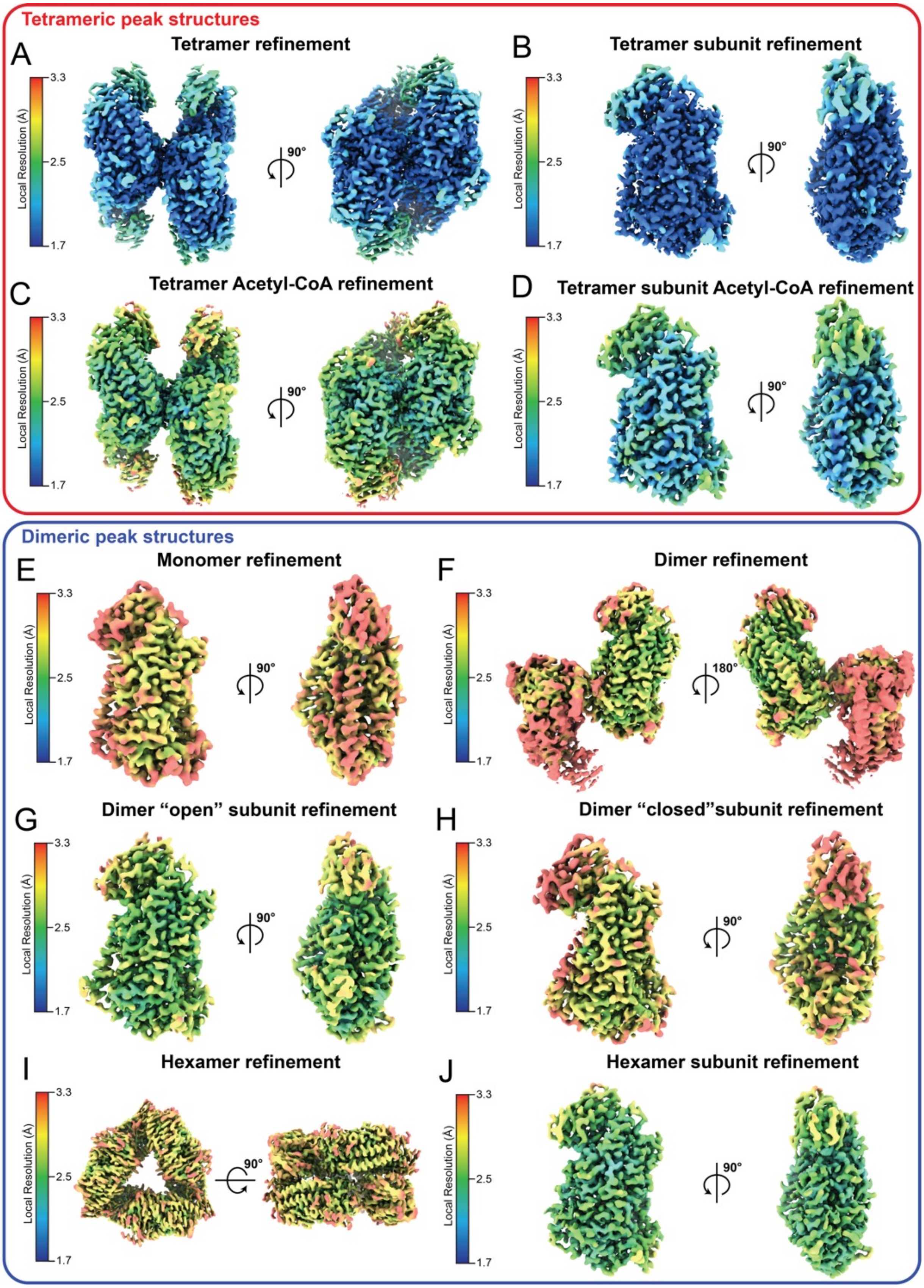


**Supplementary Figure 4. Cryo-EM density maps coloured by local resolution.** (A) Highest-resolution map of 12-LOX tetramer from the "tetrameric" SEC peak and corresponding monomer after local refinement (B). (C) The map of 12-LOX tetramer following the refinement to improve Acyl-CoA density and corresponding monomer after local refinement (D). (E) The map of 12-LOX monomer from the "dimeric" SEC peak. (F) The map of 12-LOX dimer from the "dimeric" SEC peak and corresponding "open" (G) and "closed" (H) subunits following local refinement. (I) The map of 12-LOX hexamer from the "dimeric" SEC peak and corresponding monomer after local refinement (J).

### Supplementary Figure 5

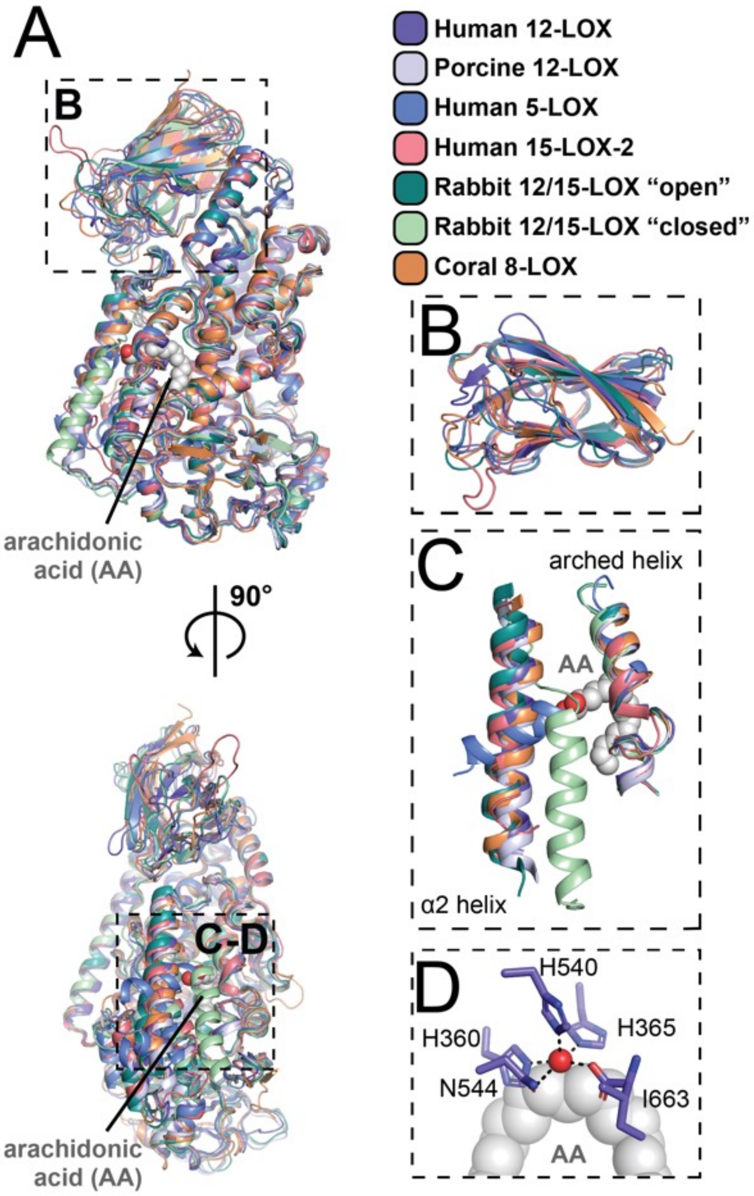


**Supplementary Figure 5. Comparison of the 12-LOX and other LOXs.** Comparison of cryo-EM structure of human 12-LOX to porcine 12-LOX (PDB: 3RDE), human 5-LOX (PDB: 3V99), human 15-LOX-2 (PDB: 4NRE), 12/15-LOX in "open" and "closed" states (PDB: 2P0M) and arachidonic acid (AA) bound Coral 8-LOX (PDB: 4QWT). (A) Overall comparison. (B) PLAT-domain comparison. (C) Arched helix and α2-helix comparison. (D) Catalytic Iron site of 12-LOX with AA displayed from Coral 8-LOX. Fe atom is shown as a red sphere. FE-coordinating residues are shown in sticks.

#### Supplementary Figure 6

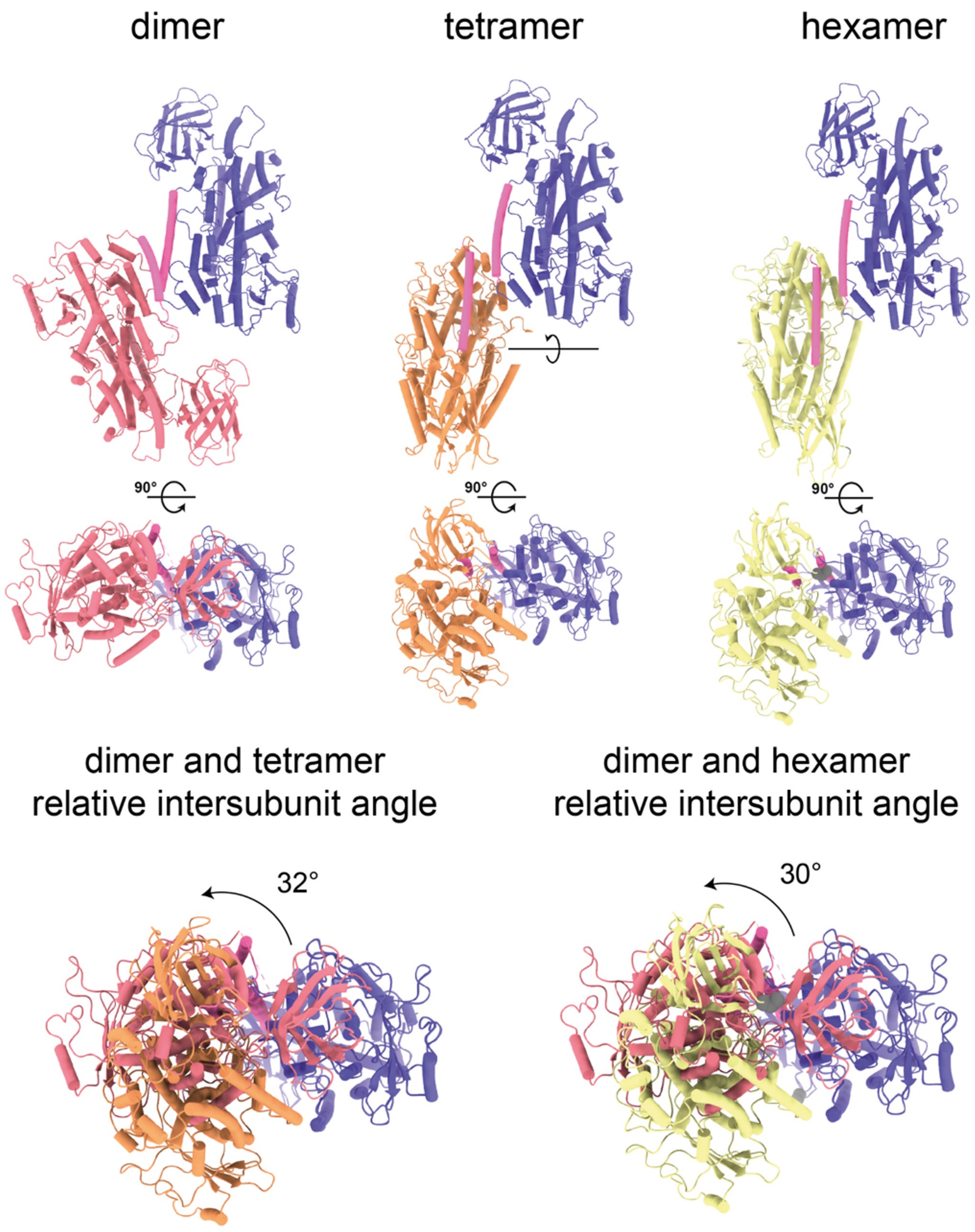


**Supplementary Figure 6. Dimers as biological units of 12-LOX.** 12-LOX exists as a dimer, dimer of dimers (tetramer), and trimer of dimers (hexamer). (Top) Dimer arrangement of each oligomeric form. (Bottom) Alignment of dimeric 12-LOX to the dimers with tetramer (left) and hexamer (right) showing the relative chain rotation within the dimers.

#### Supplementary Figure 7

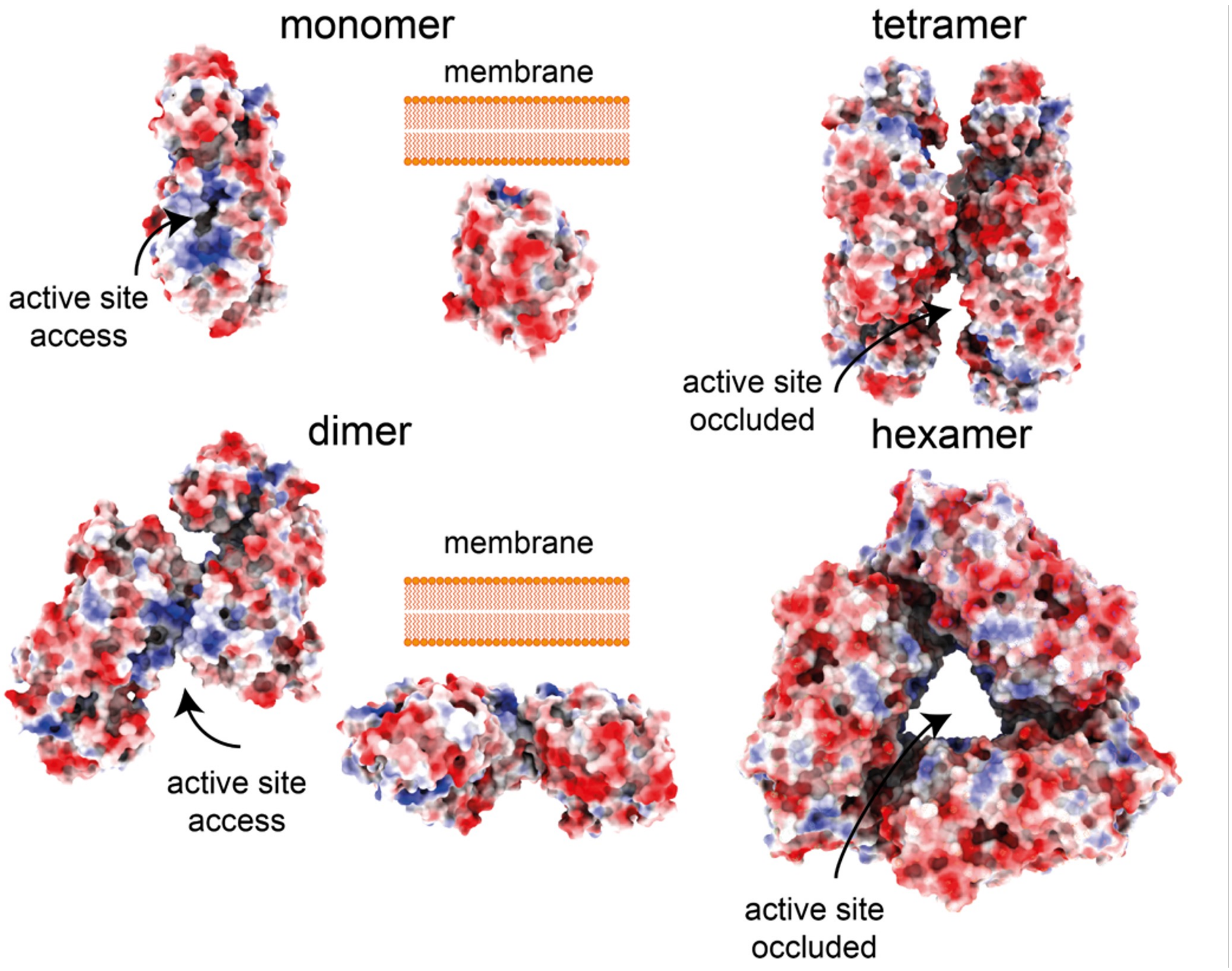


**Supplementary Figure 7. Proposed mechanism of membrane association for 12-LOX oligomers.** 12-LOX displayed as surface and coloured according to surface electrostatic potential (red is negative, white neutral and blue positive). The predicted membrane-binding surface is exposed and is available for membrane binding in 12—LOX monomers and dimers but is occluded in tetramers and hexamers.

#### Supplementary Figure 8

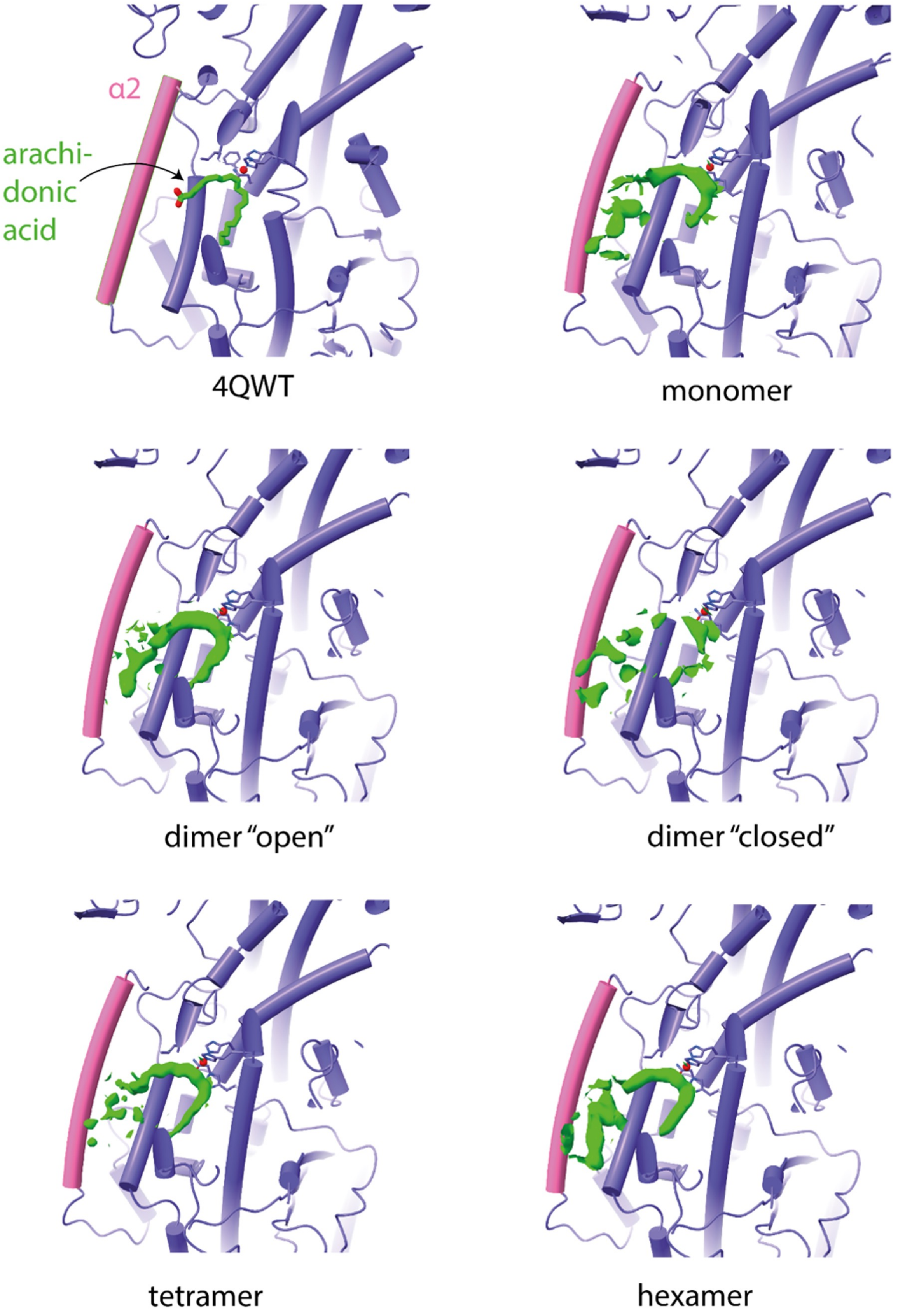


**Supplementary Figure 8. Small molecule density within the active site of 12-LOX.** In all 12-LOX subunits in the "open" conformation the active site is occupied by continuous density. 12-LOX catalytic domain is shown in purple cylindrical cartoons, the α2-helix is in pink. The density within the active site of each oligomeric form is shown as a green volume. Fe atom is shown as a red sphere.

#### Supplementary Figure 9

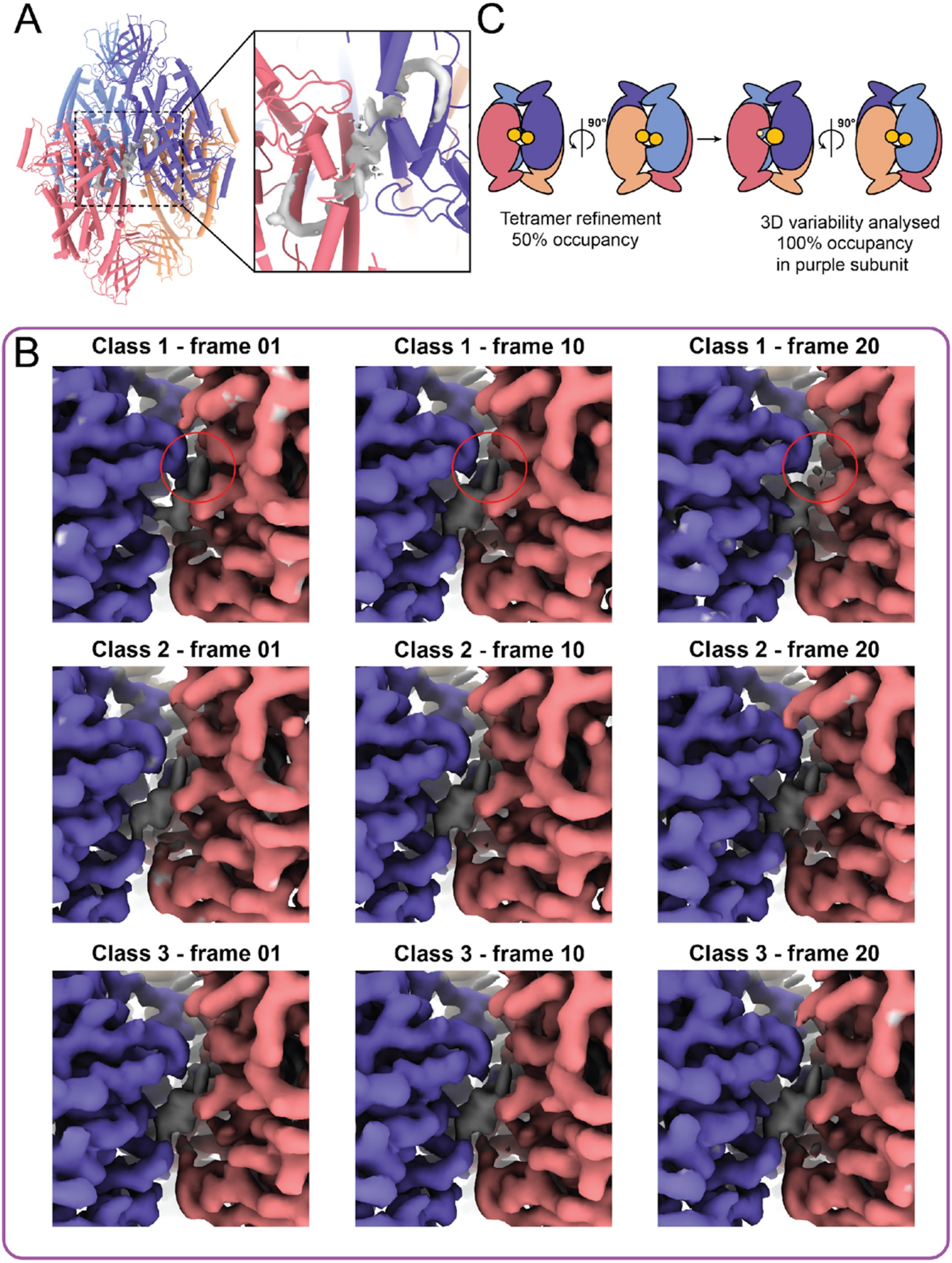


**Supplementary Figure 9. 3D-variability of the tetrameric 12-LOX showing the acyl-CoA occupancy in the active site.** (A) Model of 12-LOX tetramer with density in the catalytic site shown as grey volume. (B) 3D-variability analysis of 12-LOX tetramer. 12-LOX subunits are displayed as coloured volume (red and purple) and the catalytic site as grey volume. The red circle denotes changes observed in catalytic site density. (C) Graphical representation of ligand density.

**Table S1. Cryo-EM data collection, refinement, and validation statistics**

| | 12-LOX Monomer | 12-LOX Dimer | 12-LOX Hexamer | 12-LOX Tetramer |
|---|---|---|---|---|
| **Data Collection** | Dimer SEC peak | | | Tetramer SEC peak |
| EMD code | EMD-40039 | EMD-40040 | EMD-40041 | EMD-40042 |
| PDB code | 8GHB | 8GHC | 8GHD | 8GHE |
| Micrographs | 4182 | | | 6896 |
| Electron Dose ($e^{-}/A^{2}$) | 60 | | | 60 |
| Voltage (kV) | 300 | | | 300 |
| Pixel size (Å) | 0.82 | | | 0.82 |
| Spot Size | 5 | | | 5 |
| Exposure time | 4.8 | | | 4.8 |
| Movie frames | 60 | | | 60 |
| K3 CDS mode | yes | | | yes |
| Defocus range (µm) | 0.5-1.5 | | | 0.5-1.5 |
| **Refinement** | | | | |
| Symmetry imposed | C1 | C1 | Consensus* D3<br>1 subunit C1 | C1 (due to the asymmetry of ligand binding) |
| Particles (final map) | 35,994 | Consensus* 126,914<br>"open" 126,914<br>"closed" 34,436 | Consensus* 35,839<br>1 subunit 215,034 | Tetramer*** 311,579<br>"Oleoyl-CoA" 311,579<br>"Consensus"*455,751<br>"1 subunit" 455,751 |
| Resolution @0.143 FSC (Å) | 2.76 | Consensus* 2.53<br>"open" 2.33<br>"closed" 2.54 | Consensus* 2.63<br>1 subunit 2.24 | Tetramer***2.05<br>"Oleoyl-CoA"1.90<br>"Consensus"*1.82<br>"1 subunit"1.72 |
| $CC_{map–model}$ | 0.85 | Composite** 0.79 | Composite** 0.89 | Tetramer***0.90 |
| Map sharpening B factor ($Å^{2}$) | -30 | Consensus* -60<br>"open" -25<br>"closed" -25 | Consensus* -73<br>1 subunit -25 | Tetramer***-43<br>"Oleoyl-CoA"-39<br>"Consensus"*-47<br>"1 subunit"-44 |
| **Model Quality** | | Refined against the composite** map | Refined against the composite** map | Refined against the tetramer*** map |
| R.M.S. deviations | | | | |
| Bond length (Å) | 0.006 | 0. 004 | 0.005 | 0.004 |
| Bond angles (°) | 0.821 | 0.793 | 0.820 | 0.663 |
| Ramachandran | | | | |
| Favoured (%) | 97.25 | 97.77 | 98.16 | 98.45 |
| Outliers (%) | 2.75 | 2.23 | 1.84 | 1.55 |
| Rotamer outliers (%) | 0 | 0 | 0 | 0.04 |
| C-beta deviations (%) | 0 | 0 | 0 | 0 |
| Clashscore | 4.09 | 4.55 | 4.04 | 3.72 |
| MolProbity score | 1.33 | 1.28 | 1.19 | 1.16 |

* Consensus maps are maps prior to local refinement of individual subunits.

**For the 12-LOX dimers and hexamers we performed the final refinements against the composite maps generated in Phenix suite from local refinement maps. Dimer 12-LOX – we combined the local refinement maps for "open" and "closed" subunits. Hexamer 12-LOX – we applied the D3 symmetry to generate the entire hexamer from the locally refined single subunit.

***Tetramer 12-LOX – Following 3D variability classification of 1 subunit resulting in Oleoyl-CoA subunit map, focused refinement was performed to recover a Oleoyl-CoA bound tetramer map.

**Table S2. Enzymatic properties and inhibition of two 12-LOX peaks from SEC.** The values are from at least 3 independent experiments. The standard deviation is shown in parenthesis.

| | 12-LOX dimer SEC peak | 12-LOX tetramer SEC peak |
|---|---|---|
| **Enzymatic activity** | | |
| Kcat ($s^{-1}$) | 11.8 (0.9) | 4.8 (0.2) |
| Km ($\mu M^{-1}$ ) | 8.2 (1.0) | 3.3 (0.7) |
| Kcat/Km ($\mu M^{-1}$ $s^{-1}$) | 1.4 (0.01) | 1.4 (0.01) |
| **ML355 inhibition** | | |
| IC50 (µM) | 1.6 (0.3) | 1.4 (0.3) |
| Max inhibition (%) | 95 (5) | 92 (5) |

**Table S3. 12-LOX dimer SEC peak inhibition by acyl-CoAs.** The values are from at least 3 independent experiments. The standard deviation is shown in parenthesis.

| Substrate | IC50 ($\mu$M) |
|---|---|
| Palmitoyl-Coenzyme A (16:0) | > 200 |
| Palmitoleoyl-Coenzyme A (16:1) | > 350 |
| Stearoyl-Coenzyme A (18:0) | > 200 |
| Oleoyl-Coenzyme A (18:1) | 32 (4) |
| ɣ-Linolenoyl-Coenzyme A (18:3) | > 200 |
| Arachidonoyl-Coenzyme A (20:4) | 112 (20) |
